# Authentic field experiences for STEM teachers: Collecting Florida fossils

**DOI:** 10.1101/2020.07.02.184077

**Authors:** Bruce J. MacFadden, Jeanette Pirlo, Brian Abramowitz, Stephanie Killingsworth, Michael Ziegler

## Abstract

The extensive sedimentary sequence of Florida preserves evidence of prehistoric life spanning some 40 million years into the past. This paper describes the results of a three-pronged professional development (PD) program conducted in November 2019 in which Florida teachers from eight counties gained knowledge and experience about the process and content of paleontology at an active field research site in northern Florida. Working alongside scientists, 15 elementary, middle, and high school teachers collected 5-million-year-old fossil vertebrates and made scientifically important discoveries that advance understanding of Florida’s prehistoric life. The PD included three components: (1) a pre-trip webinar; (2) a morning tour of our museum exhibits to scaffold teacher’s understanding of the kinds of fossils they would find; and (3) the actual day-long field experience collecting fossils. A post-PD evaluation revealed gains in knowledge about, and experience with, fossils and paleontology. We also found that overall satisfaction with the PD underscored the importance of each of the three components supporting one-another—resulting in a more informative and rewarding learning experience for the participants. Fossils and the science of paleontology are a gateway for STEM learning and this subject and its extensions are applicable to existing standards in earth and life sciences at many grade levels.

## Introduction and Background

The extensive sedimentary outcrops of Florida are rich with diverse fossils that span the past 40 million years of Earth history. As the peninsula developed over time, the remains of both marine and terrestrial organisms were preserved. These fossils thereby provide evidence about ancient life, climate change, and ecosystem development during this period of time.

The Florida Museum of Natural History (FLMNH) at the University of Florida (UF) is the official repository of the state’s research collections, consisting of almost five million fossil specimens of extinct plants and animals; many of the finest examples of these are also on display in the Hall of Florida Fossils (2020). These research collections are continuously updated by fossil discoveries from Florida. Here we describe a professional development (PD) in which Florida teachers worked alongside FLMNH paleontologists at an active field research site in northern Florida. The Montbrook Fossil Site (Hulbert, 2016) has been the focus of intensive field activities since 2016. It preserves an abundant record of ancient life in an estuarine setting about 5 million years ago during the Miocene epoch. This includes extinct fishes, reptiles—including ubiquitous and diverse turtles, some birds (e.g., heron; Steadman and Tanaka, 2019), and mammals--horses, rhinos, saber-toothed cats, and a unique population of elephant-like gomphotheres, the latter of which is the basis of author Pirlo’s PhD research.

Here we describe teacher recruitment, and the Montbrook PD that included three interrelated activities, (1) a pre-trip, orientation webinar; (2) a tour of the fossil exhibits at the FLMNH prior to the field experience; and (3) the actual excavating at the fossil site. This paper also provides the results of a survey done by the teacher participants at the end of the PD. This PD experience was part of *Scientist in Every Florida School* (SEFS, 2020), a moonshot program under UF’s Thompson Earth Systems Institute.

## Conceptual Framework—Authentic STEM and Teacher Professional Development

Our Montbrook PD was informed by a conceptual framework about how to achieve effective (1) STEM learning through authentic research; and (2) best practices for teacher PD:

### 1. Authentic STEM Research Experiences

Presenting teachers with the opportunity for out-of-school experiences, such as in-field or lab based authentic science, can change how teachers view scientific content. These kinds of experiences push teachers to see new and innovating ways in which to teach students (Birmingham et al., 2018; Foster, 2018). Furthermore, Cook (2018) describes the value of placed-based geoscience field experience, like the one that we describe in this report. Studies have shown that these kinds of experiences are vital to effective teacher PD (Hall, 2004).

Silverstein et al. (2009) demonstrated that active involvement of teachers in research positively impacts students’ achievement in science. The National Science Foundation has for many years sponsored a program called “Research Experiences for Teachers” (RET, 2018), and studies have also shown that these programs and partnerships benefit both the teacher and the scientist alike (MacFadden, 2019). The main PD activity that we describe below, therefore, involves teachers collecting fossils that will subsequently be used in research; this also aligns with the importance of teachers’ understanding of the practice of paleontology. The latter is considered part of the Nature of Science (NoS), which is a core component of current K-12 standards, of which several components were addressed during our PD (Table 1).

**Table 1.** Next Generation Sunshine State Standards (NGSSS) Representative standards addressed by the professional development experience (CPALMS, 2020)

|  |  |
| --- | --- |
| Nature of Science | SC.5.N.1.2: Explain the difference between an experiment and other types of scientific investigation. |
| Life Science | SC.5.L.15.1: Describe how, when the environment changes, differences between individuals allow some plants and animals to survive and reproduce while others die or move to new locations. |
| Middle School |  |
| Nature of Science | SC.7.N.1.5: Describe the methods used in the pursuit of a scientific explanation as seen in different fields of science such as biology, geology and physics. |
| Life Science | SC.7.15.2: Explore the scientific theory of evolution by recognizing and explaining ways in which genetic variation and environmental factors contribute to evolution by natural selection and diversity of organisms. |
| High School |  |
| Nature of Science | SC.912.N.1.3: Recognize that the strength or usefulness of a scientific claim is evaluated through scientific argumentation, which depends on critical and logical thinking, and the active consideration of alternative scientific explanations to explain the data presented. |
| Life Science | SC.912.L.15.1: Explain how the scientific theory of evolution is supported by the fossil record, comparative anatomy, comparative embryology, biogeography, molecular biology, and observed evolutionary change. |
| Earth and Space Science | SC.912.E.7.3: Differentiate and describe the various interactions among Earth systems, including: atmosphere, hydrosphere, cryosphere, geosphere, and biosphere. |

### 2. Effective Teacher PD

A large body of literature exists on what constitutes effective PD, and our program described here includes many of the recommended practices (e.g., IACET, 2017): Our PD program: (1) was content focused; (2) incorporated active learning; (3) included collaboration [between scientists and teachers]; (4) used effective practices; (5) provided coaching [and mentoring]; and (6) provided opportunity for feedback and reflection. One thing that IACET (2017) recommends that we hope to do in the future is to make similar PDs of sustained duration, rather than the one-off opportunity described here.

## Teacher Recruitment

In order to recruit teachers for the Montbrook PD, we sent email notifications to school district officials across the state. We also spread further awareness and encouraged application submissions via a social media campaign through our Twitter and Facebook accounts. As a result of this campaign, we received 20 applications from teachers representing the following Florida counties: Palm Beach, Escambia, Marion, Pasco, Volusia, Broward, Alachua, and Seminole. Fortunately, were able to accommodate all who applied; of these 15 ultimately participated in the PD. Seven of the teachers had previously participated in SEFS opportunities, which helps to build a peer mentoring experience. The other eight were new to these opportunities. Consistent with our recruitment strategy, the Montbrook PD represented K-12 educators across all grade levels, including elementary, middle, and high school teachers, plus two district level STEM specialists.

## Brief Description of the Professional Development Experience

Our Florida fossil experience involved three components, all of which were planned to be mutually supportive in optimizing this PD opportunity.

1. On Monday evening, 18 November 2019, the scientists (including the SEFS coordinators) hosted a pre-trip webinar (MacFadden et al., 2019) that was enabled by Zoom (2020). The purpose of this webinar was to: (a) welcome and introduce participants, including both the teachers and the scientists; (b) describe the Montbrook Fossil Site, its context and importance; and (c) prepare the teachers for the logistics of the field experience (i.e., what to expect, what to bring). All of the teachers viewed the webinar either in real-time or subsequently via the recorded version.
2. On Saturday morning, 23 November 2019, prior to the field trip the group (teachers and scientists) assembled at the FLMNH public exhibits. Led by the scientists, they toured the Hall of Florida Fossils (2020) and Montbrook exhibit (Fig. 1) to better understand the context of the kinds and types of fossils that they would see later that day.

**Figure 1.** Montbrook fossil display at the Florida Museum of Natural History (Jeff Gage photo).
3. On Saturday, 23 November 2019, after the tour of the museum exhibits, the entire group travelled to the fossil site, which is located in Levy County about a 45-minute drive from the UF campus. We all participated in the excavations; and the teachers therefore had an authentic hands-on experience digging fossils. Each digger was assigned a 1-meter square and they spent the rest of the day carefully excavating the fossils, mostly using screwdrivers to separate these specimens from the sandy sediments (Fig. 2).

**Figure 2.** Montbrook dig on 23 November 2019. Each teacher was assigned a 1-m square and carefully excavated the fossil-bearing sediments (Jeff Gage photo).

At the conclusion of the digging in mid-afternoon, we had a brief wrap-up and reflection and then administered an IRB (Institutional Review Board) approved (UF-IRB-02, 2020), mixed methods survey consisting of quantitative questions and qualitative (open-ended) questions. All of the teacher participants (N=15) completed the survey.

## Results and Findings

### Fossil Collecting

During the day’s excavations more than 100 fossils were found, including scientifically important specimens like a premolar of *Nannipus aztecus* a small, three-toed horse. The molar was found by Escambia County 5^th^grade teacher Rosalyn Rohling (Lindsay, 2019). This tooth is important, not just because it is part of the representative fauna at the site, but also because its shape indicates the diet of this species. By examining the grinding or occlusal surface (Fig. 3), as well as the crown height of the tooth, paleontologists can infer the type of vegetation this ancient horse ate; it was probably a grazer (also see lesson plan on this topic described in Bokor et al., 2016).

**Fig. 3.** Left: Escambia County educators (left to right): Carol Myers, Suzanne Delay, Rosalyn Rohling, and Kathleen Stanhope. Rosalyn collected, and is holding, a tooth of the tiny extinct horse, *Nannippus aztecus*, from Montbrook, much to the delight of all of the participants. Right: Close up view of the upper tooth showing the grinding (occlusal) surface (Victor Perez photo).

Most of the participants found fossils, including the ubiquitous turtle shell pieces from *Trachemys inflata*, an extinct relative of the common Red-Eared Sliders of Florida. There were also many fragments of the freshwater fishes, including vertebrae, gar scales, and spines. All of these fossils tell an important story--they are incredibly informative of the environment, and together with the horse tooth, they help site paleontologists better understand the ancient ecology of northern Florida 5 million years ago.

## Survey Results and Feedback

While designing the PD, we identified two goals that we hoped our participants would achieve by its completion: (1) To experience authentic field research through the collection of fossils; and (2) increase understanding of the process of paleontology from the field to the exhibit display.

As expected, goal 1 was easily achievable because participants were required to spend time in the field collecting fossils, but goal 2 really depended on the interactions between the scientists and the teachers. Based on the survey results, the majority of our participants felt that they had increased their understanding of the process of paleontology and that their confidence to do science had increased as a result of participating in the PD (Fig. 4). Our hope was that the webinar would be viewed as an important component to the experience, but we did not expect the very positive response that it received: The majority of participants identified the webinar as a “way to set the stage” and to “get [them] excited” for the experience because it provided an opportunity to “set expectations” for the field work.

**Fig 4.**
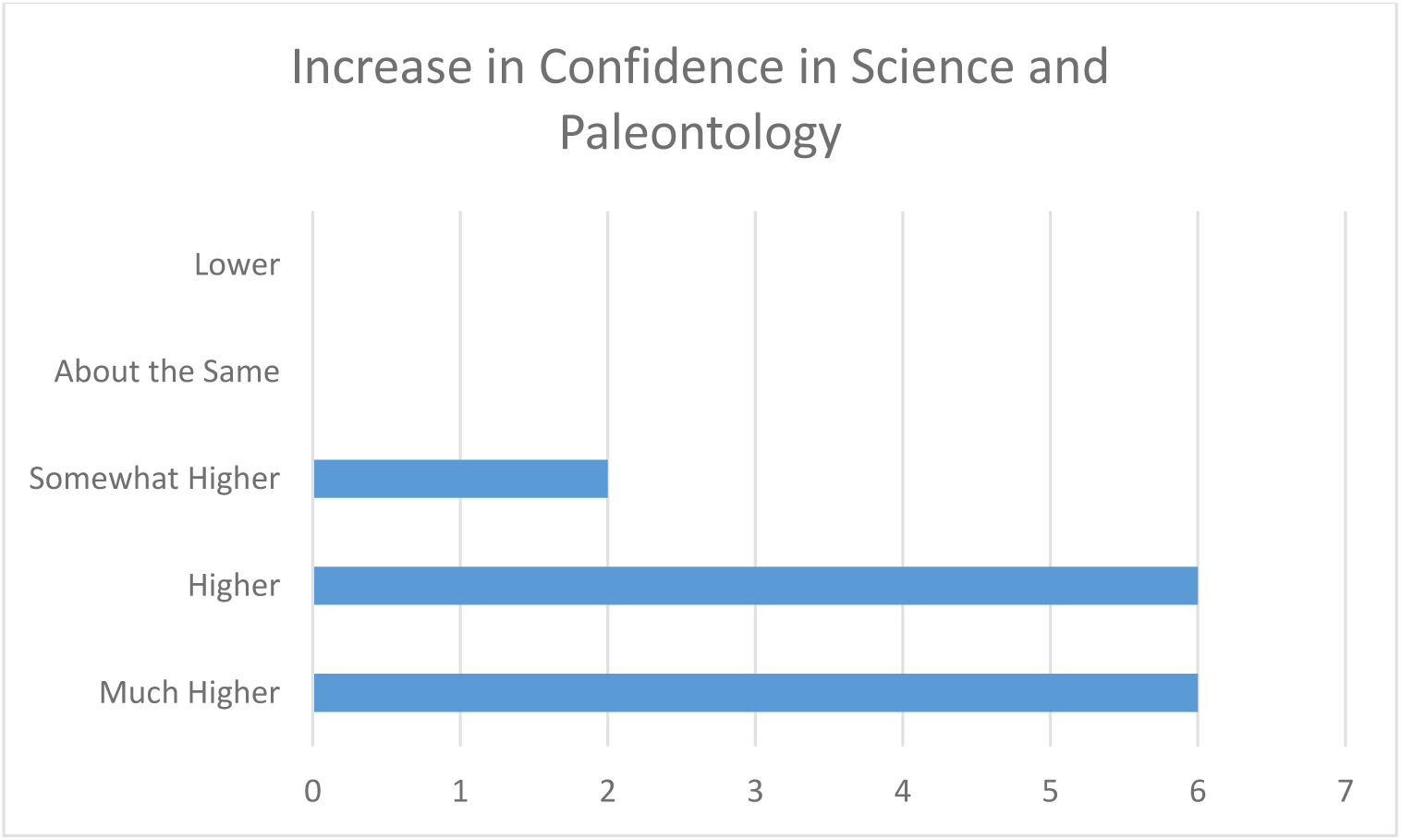
Graph showing participants’ responses (14 of 19) when prompted to rate “how does your confidence to do science and/or paleontology NOW compare to PRIOR to the PD?”

Participants had an opportunity to describe how they would use their experience back in the classroom. Many described the importance of sharing the experience through photos and stories with their students. One participant described the learning experienced: “I feel much more confident talking about limestone, estuarine environments, [and] paleo animals that inhabited Florida. I will integrate into [the] Florida History and Science Standards.” While another expressed their own discoveries: “First, I was unaware that Florida had any dig sites. Now I know about a site and how to dig. It was so amazing. I will be sharing this for years to come.” One participant had “[made] a video of the trip to show my peer teachers and six classes.” Based on the survey results, it appears that we achieved our goals and provided an educational and effective PD for these teachers.

## Wrap-Up and Take-Away Messages

Overall, the Montbrook PD, consisting of three interconnected components (webinar, museum, and field), achieved our programmatic goals, and as indicated by the teachers’ surveys, their needs as well. Given previous field work, the excavation of so many fossils was expected, and added to the research collections at the FLMNH. The discovery of the rare, tiny horse tooth described above was celebrated by the participants at the field site. This discovery was also subsequently promoted in the popular media that reached the local TV audience in Escambia County and via Twitter (Lindsay, 2019), where it reached 782 followers (Engagement 29%) and on Twitter had 1.022 Impressions (Engagement 5.8%).

Any PD like this one can be improved upon, and two points come to mind: (1) Our PD should also intentionally have a fourth component, i.e., building a lasting specific connection/partnership with individual teachers and scientists, which is also part of the mission of SEFS. In so doing, they could further extend this experience into the classroom via scientists’ role model visits. (2) If we had another half day, we could take the teachers back into the museum collections behind-the-scenes, and in so doing, allow them to understand the entire practice of paleontology, ranging from field discovery, to curation back in the lab, to developing museum displays, from which the public can learn.

The Montbrook Fossil Site remains an important field research site to advance knowledge about Florida’s prehistoric life. It also will continue to serve as the research emphasis of UF scientists, including students, staff, and faculty. We expect to hold additional PD’s in the future that are focused on Montbrook for as long as it remains an active field site. We therefore hope that we can continue to engage Florida teachers in the enterprise of authentic research experiences.

**Fig. 5.** Group photo of teachers and scientists from the Montbrook Fossil Site, 23 November 2019 (Jeff Gage photo).

## Acknowledgements

We thank the participants for their input and enthusiasm, Rebecca Burton for help with the social media analytics, and Jeff Gage and Victor Perez for taking the photos. This PD is part of the *Scientist in Every Florida School* Moonshot project funded by the UF Office of the President and Provost. The survey was approved (exempt) by UF IRB 201903056. This project is also part of a National Science Foundation Graduate Research Fellowship (GRFP) Broader Impacts plan for Jeanette Pirlo.

